# Deciphering a beetle clock: individual and sex-dependent variation in daily activity patterns

**DOI:** 10.1101/2024.03.18.585527

**Authors:** R Reshma, Tobias Prüser, Nora K. E. Schulz, Paula M. F. Mayer, Maite Ogueta, Ralf Stanewsky, Joachim Kurtz

**Author notes:** Corresponding author: Prof. Dr. Joachim Kurtz, Hüfferstr. 1, 48149 Münster. Shared senior author (Ralf Stanewsky, Joachim Kurtz).

## Abstract

Circadian clocks are inherent to most organisms, including cryptozoic animals that seldom encounter direct light, and regulate their daily activity cycles. A conserved suite of clock genes underpins these rhythms. In this study, we explore the circadian behaviors of the red flour beetle *Tribolium castaneum*, a significant pest impacting stored grain globally. We report on how daily light and temperature cues synchronize distinct activity patterns in these beetles, characterized by reduced morning activity and increased evening activity, anticipating the respective environmental transitions. Although less robust, rhythmicity in locomotor activity is maintained in constant dark and constant light conditions. Notably, we observed more robust rhythmic behaviors in males than females with individual variations exceeding those previously reported for other insect species. RNA interference targeting the *Clock* gene disrupted locomotor activity rhythms. Our findings demonstrate the existence of a circadian clock and of clock-controlled behaviors in *T. castaneum*. Furthermore, they highlight substantial individual differences in circadian activity, laying the groundwork for future research on the relevance of individual variation in circadian rhythms in an ecological and evolutionary context.

## Introduction

Virtually all lifeforms from bacteria to humans express daily patterns in their physiological and behavioral processes, known as circadian rhythms, which enable them to accommodate the recurring changes in their environment (Saunders, 2002). These rhythms are not only reactions to the environment but are endogenously controlled by molecular pacemaker systems called circadian clocks, which can be entrained to the external periodic cues such as light and temperature, which are collectively known as zeitgebers (Bell-Pedersen et al., 2005). Coordinating activities in anticipation to the periodic environmental changes has obvious adaptive functions such as escaping adverse environmental conditions, ensuring proper availability of food and mates, synchronizing behaviors with conspecifics and reducing competition or predation risks (Favreau et al., 2009; Vaze and Sharma, 2013). Apart from these extrinsic advantages, a circadian clock confers an intrinsic advantage of maintaining a stable and optimal temporal compartmentalization of biological processes regardless of any irregular fluctuations in the environment. Even animals living in aperiodic conditions are known to have a functioning molecular clock (Trajano and Menna-Barreto, 1996) which points towards its intrinsic adaptive values.

Circadian clocks operate similarly across different taxa. Their cellular mechanism is comprised of transcription-translation feedback loops of clock genes and their protein products, capable of producing self-sustained circadian oscillations (Doležel, 2023; Helfrich-Förster, 2004). Detailed knowledge about the molecular mechanisms of animal clocks is available mainly from the studies in mouse and fruit fly as representatives of mammals and insects, respectively. In the main circadian feedback loop of fruit flies, the clock proteins PERIOD (PER) and TIMELESS (TIM) represent the transcriptional repressors, whereas in mammals CRYPTOCHOROMES (CRY1 and CRY2) dimerize with the PERIOD proteins (mPER1, mPER2 and mPER3) to perform the equivalent role. This transcriptional inhibition occurs by blocking of their transcriptional activators CLOCK and CYCLE (BMAL1 is the mouse homolog of CYCLE). A striking difference between the mammalian and *Drosophila* clock lies in the mechanism by which light resets and synchronizes the clock with the environment. Light induces TIM degradation in flies whereas it leads to the transcriptional activation of the *Period* genes in mice (Helfrich-Förster, 2004).

Insect clocks are best documented in *Drosophila*. However, given the enormous diversity of insect species and their habitats, it is plausible that their molecular clock mechanisms have diversified in response to their specific environmental conditions (Kotwica-Rolinska et al., 2022; Tomioka and Matsumoto, 2015). Diversities in some clock components are already known, for instance, certain insects like beetles and honeybees possess the mammalian light-insensitive cryptochrome, CRY2, unlike *Drosophila*, which possesses only the photosensitive CRY1. On the other hand, mosquitoes and monarch butterflies are known to possess both CRY1 and CRY2 (Yuan et al., 2007). This underscores the significance of investigating a broad range of insect models for gaining a comprehensive understanding of various aspects of the circadian clock.

*Tribolium castaneum,* the red flour beetle, is a widely found pest of stored grains and an important model organism of increasing relevance in many fields such as classical genetics, population genetics, insect development and epigenetics (Campbell et al., 2022). These flour beetles are cryptozoic, meaning that they are animals living mostly in darkness (Brown et al., 2009) They have very little exposure to light in their habitat of stored grains or behind tree barks, their supposed original environment (Good, 1936; Sokoloff, 1974). Their cryptozoic adaptations include loss of photoreceptors sensitive to blue light (Jackowska et al., 2007) and an increase in odorant and gustatory receptors (Richards et al., 2008). Even though the adults are normally known for being negatively phototactic (Park, 1934), they crawl to flour surfaces from time to time where they can get direct light exposure as observed in laboratory populations (Misra and Englert, 1985). Copulation usually occurs on the flour surface as well (Park, 1934). They are also likely to experience temperature variations in their natural habitats. Despite spending most of their life in the absence of light, we know that they still possess the genes of a functional molecular clock (Li et al., 2018). However, whether *T. castaneum* indeed displays rhythmic activity profiles, driven by a functional circadian clock, has not been convincingly demonstrated.

*T. castaneum* has homologs for all clock genes, known in *D. melanogaster* (Richards et al., 2008). First studies on the molecular clock in *T. castaneum* revealed that *Clock* (*Clk*), *cycle* (*cyc*), *period* (*per*) and *timeless* (*tim*) are all rhythmically expressed (Li et al., 2018). Instead of the *Drosophila*-type cryptochrome, CRY1, these beetles possess the light-insensitive CRY2, the function of which remains unclear. It has been suggested that CRY2 plays a role in inhibiting CLK and CYC, thus acting as a negative element of the feedback loop as known in other insects such as Hymenoptera and Heteroptera (Kotwica-Rolinska et al., 2022; Yuan et al., 2007). However, *T. castaneum* also possesses the *Drosophila*-type TIM (Kotwica-Rolinska et al., 2022) along with the mammalian CRY2 making it a unique model to study insect clocks, also possibly in an evolutionary context of transition from the *Drosophila*-type to mammalian-type clocks.

In insects, functional circadian clocks, and the rhythms they produce are traditionally evaluated by measuring an individual’s variation in locomotor activity during the course of the day (Chiu et al., 2010). It has been shown that *T.castaneum* exhibits rhythmic locomotor activity under light dark cycles (LD; Rath et al., 2021) as well as in the absence of external cues in constant darkness (DD; Abe et al., 2021). However, both studies report only average data of individually kept beetles, without determining the overall rhythmicity (i.e., how many beetles behaved rhythmically in a certain condition), nor showing individual actograms, which would allow to estimate the individual variation within the population. It is important to investigate these behavioral variations as an organism reacts to and modifies its environment through its behavior (Duckworth, 2009) and therefore having sufficient behavioral variation can readily help populations to adapt faster to changing environments (Bolnick et al., 2003). Furthermore, to our knowledge, there are also no studies exploring whether the circadian clock in these beetles can be synchronized by temperature fluctuations.

In the present study, we monitored the locomotor activity patterns of *T. castaneum* to characterize in detail the circadian profiles in their movements. We studied light as well as temperature entrainment and the effect of constant conditions on an individual level to characterise the behavioral variations existing within the population. We also performed RNAi to study the changes in activity patterns upon knockdown of the clock gene encoding the key circadian transcription factor CLK. As *T. castaneum* is an important model organism, knowledge about their biological clock and how it influences their behavior and physiology can add to the knowledge in their proper laboratory rearing conditions and proper timing of experiments. It can serve as an exceptional model system for investigating the molecular clock mechanisms of Coleoptera, where our current understanding is comparatively limited and thereby also advancing our overall comprehension of insect clocks.

## Materials and Methods

### Beetle cultures

For this study, the Croatia 1 (Cro1) strain, collected in Croatia in 2010 and a commonly used lab-adapted strain San Bernardino (SB), from San Bernadino, California, USA, established by Sokoloff in 1960’s were used (Milutinović et al., 2013). The populations were kept at standard laboratory conditions of 30 °C, 70% humidity and a 12 h:12 h light: dark cycle (lights on at 08:00 and lights off at 20:00; light from flourescent bulbs, 4000 K, cool white, 3350 lumen) in plastic boxes or glass jars with foam stoppers to allow air circulation. The standard beetle diet consisted of heat-sterilized (75 °C for at least 24 h) organic wheat flour (Bio Weizenmehl Type 550, dm - Drogerie Market) mixed with 5% brewer’s yeast powder (referred to as “flour” from now on).

### Locomotor activity

To monitor locomotor activity patterns, we used Drosophila Activity Monitoring System (DAM5H-4, Trikinetics Inc., Waltham, MA, USA; referred to as “DAM5 system” in the following). Individual beetle’s movements interrupt one of four infrared light beams intersecting the glass tubes housing single beetles, at equal distances from each other (Rosato and Kyriacou, 2006, Fig S1) which enables the recording of an individual’s movements. Sexually matured virgin beetles of known sex were placed into an individual glass tube (diameter: 5mm, length:65mm) with a flour-disc (40μl of flour/water solution (0.15g/ml) dried overnight) as food inside on one end and sealed with cotton plugs on both ends. The glass tubes were assembled into the DAM5 systems which further were placed into incubators for the recording of activity. The standard rearing conditions were maintained in the incubators throughout the period of recording unless stated otherwise. Before the first day of recording, beetles acclimatized for at least six hours in the activity monitors.

### Circadian activity patterns under light entrainment followed by constant darkness (LD **DD)**

For the general characterization of the circadian activity patterns under 12:12 Light/Dark (LD) and subsequent constant darkness (DD), we used the Cro1 population (the beetles used belong to the handling control of a knockdown experiment). After sex discrimination we individualized the pupae in wells of a 96-well flat-bottom micro reaction plate (Type F, Sarstedt) containing 0.08 g of flour. 32 adults per sex, which were at least two-week-old post eclosion were used for monitoring locomotor activity for an initial seven days during LD at 30°C (normal rearing temperature) at a light intensity of 20 lux followed by seven days of DD.

### Circadian activity patterns in constant light after light entrainment (LD LL)

To study the effect of constant light on the circadian activity patterns, we monitored beetles of the Cro1 population in constant light (LL) after entrainment to LD, at two different light intensities: 20 lux and 100 lux. Around two weeks post eclosion, we used 32 adults per sex, which were individualized since sex discrimination at pupal stage, for each light intensity. We recorded their activity for an initial seven days in the LD followed by six days in LL.

### Comparison of activity profiles between two laboratory populations

For comparing the circadian activity patterns of the two laboratory populations Cro1 and SB, we separated the pupae according to sex and kept them in groups of 24 individuals in petri dishes (∅ 10 cm, Waldeck) with *ad libitum* flour. For each population, 24 adults per sex which were at least 11 days post eclosion (SB population develops slower than Cro1) were used. The activity recording was carried out in LD (20 lux) for 7 days followed by another 7 days of DD. During LD, a technical issue caused the temperature and humidity to drop from 30°C to 25°C and from 70% to 50%, respectively for 8 hours (from midnight until lights on at 8am).

### Circadian activity patterns under temperature entrainment

To test if daily temperature cycles can entrain the locomotor activity patterns in variations.

### Activity profiles after knockdown of *Clk*

To test, if rhythmic behavior depends on the canonical molecular clock mechanism, we performed RNAi-knockdown (Tomoyasu et al., 2008) of the key circadian clock gene *Clk* at the pupal stage using the Cro1 population. After sex discrimination, pupae were glued with their last abdominal segment onto a glass slide for the dsRNA injection. The pupae were injected between the second and third lowest segment of their abdomen with dsRNA of either *Clk* or jellyfish eGFP , which served as the control for the dsRNA injection treatment (Schulz et al., 2022). The dsRNA construct of eGFP was produced in our lab (via cloning followed by PCR and *in vitro* transcription using the T7 MEGAscript Kit (Ambion by Life TechnologiesTM GmbH, Darmstadt, Gemany)). The dsRNA construct for *Clk* was obtained from EupheriaBiotech (Dresden, Germany) and has previously been used in the ibeetle RNAi screen (Schmitt-Engel et al., 2015, Table S1). A third treatment group served as the pure handling control where the pupae glued on to the glass slides did not receive any injection. dsRNA was injected at a concentration of 1000 ng/μL dissolved in Phosphate-buffered saline (PBS) using a microliter injector (FemtoJet, Eppendorf AG, Hamburg, Germany) and borosilicate glass capillaries (100 mm length, 1.0 mm outside diameter, 0.021 mm wall thickness; Hilgenberg GmbH, Malsfeld, Deutschland). Upon eclosion, the beetles were kept individually in 96-well plates with 0.08 g flour until the start of the activity measurements in the DAM5 system.

The activity measurements started two weeks after eclosion. The activity of beetles from all three groups were monitored in LD (light intensity of 100 lux) for seven days followed by seven days in DD.

To confirm the successful downregulation following the RNAi treatment, we measured *Clk* expression by qPCR. Fourteen days after eclosion, we shock froze the individuals in liquid nitrogen. Furthermore, we usually pooled 3-4 whole bodies per replicate. All samples were stored at -80°C until the RNA extraction.

We extracted total RNA by combining a TRIZOL-chloroform approach with a spin column kit (SV Total RNA Isolation System, Promega (Peuß et al., 2016)). In short, we ground the TRIZOL-submerged samples with a sterile pestle until no larger tissue parts were visible. Samples then went into an ultrasound bath to further ensure disintegration of tissue. After ten minutes of incubation, samples were spun down and the supernatant was mixed with chloroform. Another centrifugation step ensured formation of phase separation. We then transferred the upper aqueous phase mixed with lysis buffer onto the spin column and proceeded according to manufacturer’s protocol. Finally, RNA was eluted with 80µl nuclease free water and stored at -80°C until further use.

For the qPCR, we first reverse transcribed RNA to cDNA using the RevertAid First Strand cDNA Synthesis Kit (Thermo Scientific) and the provided oligo(dT)_18_ primer, following the manufacturer’s instructions. Before the cDNA was used in the qPCR it was diluted 1:5 with nuclease free water.

We performed qPCRs with technical duplicates and two housekeeping genes *rp49 and RpL13 a* to normalize target gene expression. Amplification efficiency of primers was previously established (Table S2) as well as the uniform expression of the housekeeping genes across sexes and treatments. The qPCR ran on a LightCycler 480 System (Roche) with KAPA SYBR Fast mastermix (Sigma-Aldrich).

For each RNAi treatment and control, we measured gene expression separately for males and females in four biological replicates. We included samples in the downstream analysis if the standard deviation between the technical replicates remained below 0.5 and the melting curves showed only one clear peak.

### Behavior analyses

After two weeks of recording, we extracted the raw data of locomotor activity in activity bins of 30 minutes using DAMFileScan software (TriKinetics). We looked for activity patterns from the average histograms during different entrainment and constant conditions, and differences in average activity levels, and proportion of rhythmic beetles using data from individual beetles.

For the average comparison of diel patterns in movement, average histograms of mean activity in 30-minute intervals for all individuals belonging to the same treatment and sex group across the days of the same environmental condition, were generated using the Fly-toolbox (Levine et al., 2002b).

Individual beetles were scored to be rhythmic or arrhythmic based on a combination of qualitative and quantitative measures using double plotted actograms, autocorrelation (Levine et al., 2002b) and χ -periodograms (Sokolove and Bushell, 1978) generated using Fly-toolbox in MATLAB (MATLAB and Statistics Toolbox Release 2021b). Double plotted actograms were used for the visual analysis to identify rhythmic patterns. Each row in a double plotted actogram displays the activity in two consecutive days for which the activity of the second day is replotted in the left part of the next row to inspect for the activity patterns horizontally and vertically. A beetle is categorized as rhythmic if activity patterns are repeated for at least half of the days of activity recording. To reduce the subjectivity from the visual detection of rhythmicity, quantitative thresholds from autocorrelation and periodograms were also used in combination with the visual inspection of actograms (Levine et al., 2002b).

Autocorrelation graphs identify patterns in the data by examining the correlation between the movement signal and a time-shifted version of itself. This analysis helps detect both the strength of the rhythm (RS-value) and the duration of the identified period (p). The χ2-periodogram assesses rhythmicity by testing various potential periods within the data and evaluating their fit, represented by the Qp value. The dominant signal period is determined based on the highest Qp value, while a 95% confidence interval is used to evaluate the significance of the chosen period. Additionally, the h-value indicates to which extend the Qp value of the selected period exceeds this confidence interval. A beetle was categorized as rhythmic only if it met all three criteria: having a clear rhythmic pattern in the actogram, RS>1 in the autocorrelation and h > 10 in the periodogram.

The circular phase plots showing the phase of individual beetles (Levine et al., 2002b) were also plotted using MATLAB. For the analysis of activity levels, activity in 1-minute intervals were averaged over all monitoring days of a specific condition for individual beetles. The bar graphs, plotting proportion of rhythmic beetles and boxplots for the average activity were generated using ggplot2 (Wickham, 2016) in RStudio version 2021.09.1 (RStudio Team, 2021) under R version 4.1.2 (R Core Team, 2021). We did some final editing of supplementary figure panels using Inkscape (Developers, Inkscape Website, 2020)

## Statistics

All the statistical analysis for the rhythmicity and average activity levels were The analysis for rhythmicity was done on binary data indicating either presence and activity levels were analyzed in generalized linear mixed effect models (GLMMs) (Table S3) with the according error distributions. The final models were derived after p-value based stepwise model simplification.

For the statistical analysis of gene expression after RNAi treatment, we used the between the treatment groups.

## Results

To foster our understanding of the workings of the molecular clock in flour beetles and to further characterize the circadian patterns in their behavior, we first monitored and analysed the locomotor activity rhythms of an outbred population (Cro1) of *T. castaneum.* We tested the beetles’ behavior under different light and temperature cycles for both sexes. Additionally, we compared the outbred strain (Cro1) to an inbred strain (SB), which has adapted to laboratory conditions for decades, to infer strain specific behavioral differences. Finally, we downregulated *Clk* gene expression via RNAi to investigate the role of this key circadian clock gene in generating the behavioral patterns.

### Locomotor activity patterns under light entrainment

We monitored the locomotor activity patterns of beetles from the Cro1 population under LD (20 lux) conditions followed by DD. The average activity profiles of beetles showed a clear and repeatable diurnal rhythm (Fig 1 a-d). The daily average activity patterns were characterized by a consistent reduction in activity levels just before the onset of morning light in all the experiments, which we refer to as ‘morning dip’ (Figure 1 a). They reacted with a startle response of increased activity observed in response to the onset of the morning light stimulus (Figure 1a). This response fades quickly within 30 minutes and is not observed under DD conditions. Upon more natural like conditions with a linear ramping, extending the lights-on transition to 2 hours, the startle response either disappeared or appeared at the end of the ramping period, but as a lower peak compared to the non-ramping conditions (Fig S2). Further, beetles gradually increased their activity levels throughout the late afternoon, a behavior which we accordingly (and in analogy to a similar pattern observed in *D. melanogaster* (Hamblen-Coyle et al., 1992)) call evening anticipation (Figure 1a). With the start of the dark phase in LD, the beetles slightly reduced their activity levels from their highest peak during the evening anticipation but remained highly active throughout the night until they reduced their activity considerably during the morning dip (Figure 1 a and c).

**Figure 1:**
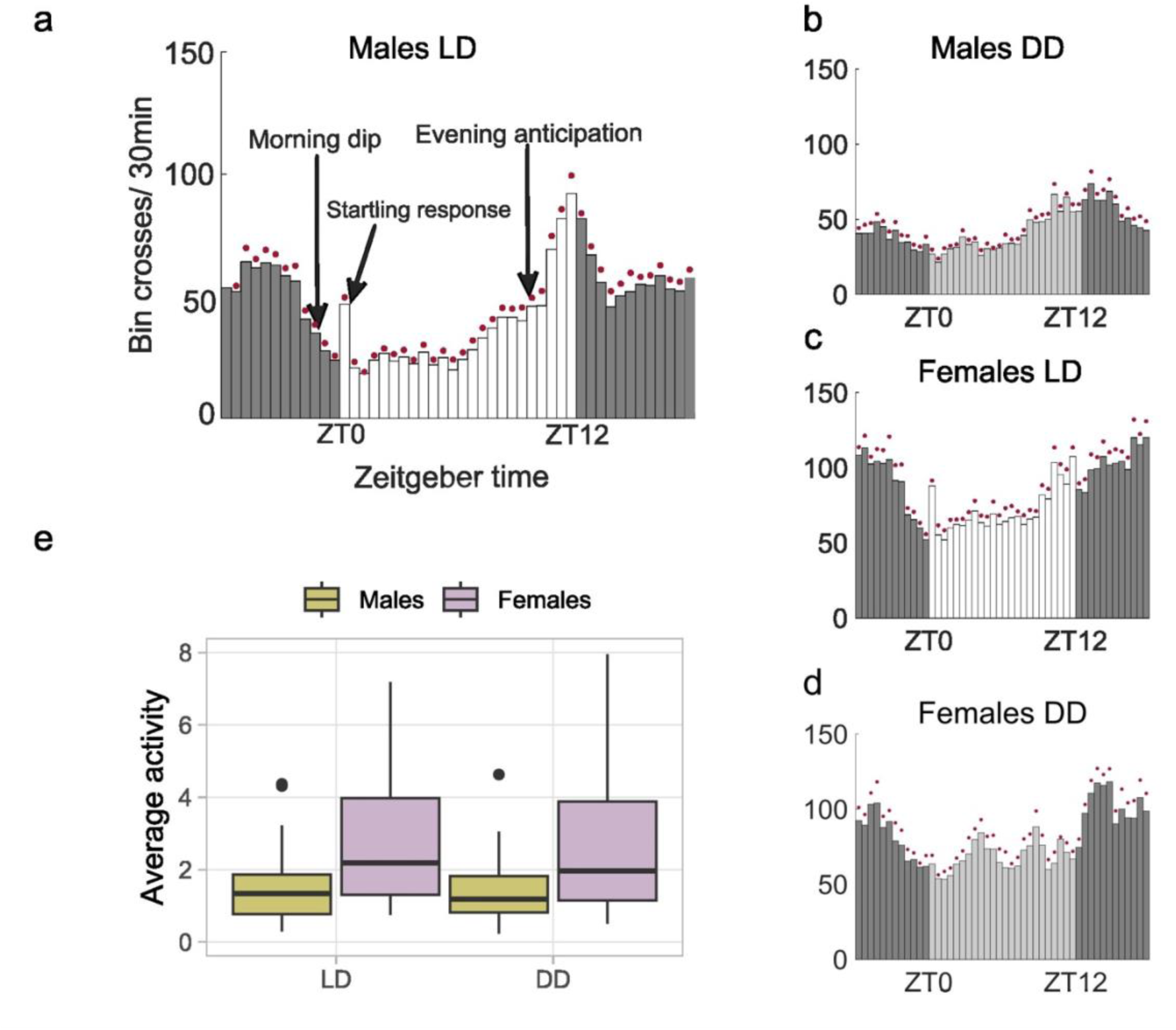
Locomotor activity patterns of *T. castaneum*. a-d: Average histograms showing mean activity of Cro1 beetles over the 7 days of recording in 30-minute bins under LD (light entrainment of 20 lux) and DD; e: Boxplots showing activity in 1-minute bin for individual beetles averaged over 7 days in LD and DD (males: n = 31, females: n=32).

### Locomotor activity patterns under constant darkness

In DD, the beetles’ average activity profiles mirrored those observed during the LD except for the obvious absence of the startle response. Before the start of the subjective morning, there was a clear reduction of activity similar to the morning dip and a gradual increase of activity throughout the subjective late afternoon that resembled the evening anticipation (Fig 1 b and d). In addition to the generally higher levels of activity observed in females compared to males, the average histograms also illustrate the irregular and noisy patterns exhibited by the females (Fig 1 a-d). The analysis of individual average activity in 1-minute bins also reveals a significantly lower average activity for males (GLMM, estimate = -0.588, p < 0.001, Fig 1e). In terms of individual rhythmicity, 52% of males and 22% of females showed rhythmic patterns in LD whereas the proportion of rhythmic beetles was similar (32% for males and 28% for females) in DD conditions (Table 1). However, upon statistical analysis of individual rhythmicity, there was no significant effect for the sex (GLMM, estimate = 0.338 p = 0.665) or the light condition (GLMM, estimate = -0.491 p=0.489) whereas there was a weak trend for the interaction between the effects of sex and light condition (GLMM, estimate = 1.727, p = 0.089)

**Table 1:** Proportion of rhythmic beetles in different experiments

| Experiment |  | Percentage of rhythmic beetles |  |  |  |
| --- | --- | --- | --- | --- | --- |
|  |  | Entrainment |  | Constant condition |  |
|  |  | Males | Females | Males | Females |
| LD (20 lux)/DD |  | 51.6% | 21.9% | 32.3% | 28.1% |
| LD LL | 20 lux | 50.0% | 18.8% | 18.8% | 9.4% |
|  | 100 lux | 75.0% | 31.3% | 12.5% | 3.1% |
| Population difference in LD DD | Cro 1 | 52.2% | 29.2% | 5.0% | 5.3% |
|  | SB | 58.3% | 56.5% | 30.0% | 27.3% |
| Temperature entrainment |  | 45.8% | 25.0% | 45.8% | 16.7% |
| Temperature entrainment with ramping |  | 35.4% | 33.3% | 28.1% | 21.9% |
| Knockdown of *Clk* | Naïve | 45.8% | 20.8% | 16.7% | 12.5% |
|  | RNAi control | 58.3% | 33.3% | 17.4% | 29.2% |
|  | *Clk* knockdown | 33.3% | 16.7% | 8.3% | 4.2% |

### Variations in locomotor activity patterns

A substantial amount of individual variation can be seen in the rhythmic patterns in LD (Figure 2 a-f). Among the rhythmic beetles, not all show identical patterns. Activity peaks of many beetles start just before the lights turn off, and among these, some have their peak of activity for a short duration before the lights turn off (Fig 2 c) or extending further throughout the dark phase of the LD (Fig 2 a). The individual variations in phase of their activity can also be seen from the circular phase plot (Fig 2 g). The females have more coherent phase compared to the males. There was variation in terms of whether the beetles could consistently maintain the rhythmicity throughout both light conditions as well (Fig 3 a-d). Not all beetles that were rhythmic in LD were able to maintain their rhythmicity during DD. 9 out of 31 males and 2 out of 32 females were rhythmic in LD but showed arrhythmic behavior in DD. There was a small proportion of beetles (3 out of 31 for males and 4 out of 32 for females) that showed rhythmic behavior only in DD (Fig 3c, e).

**Figure 2:**
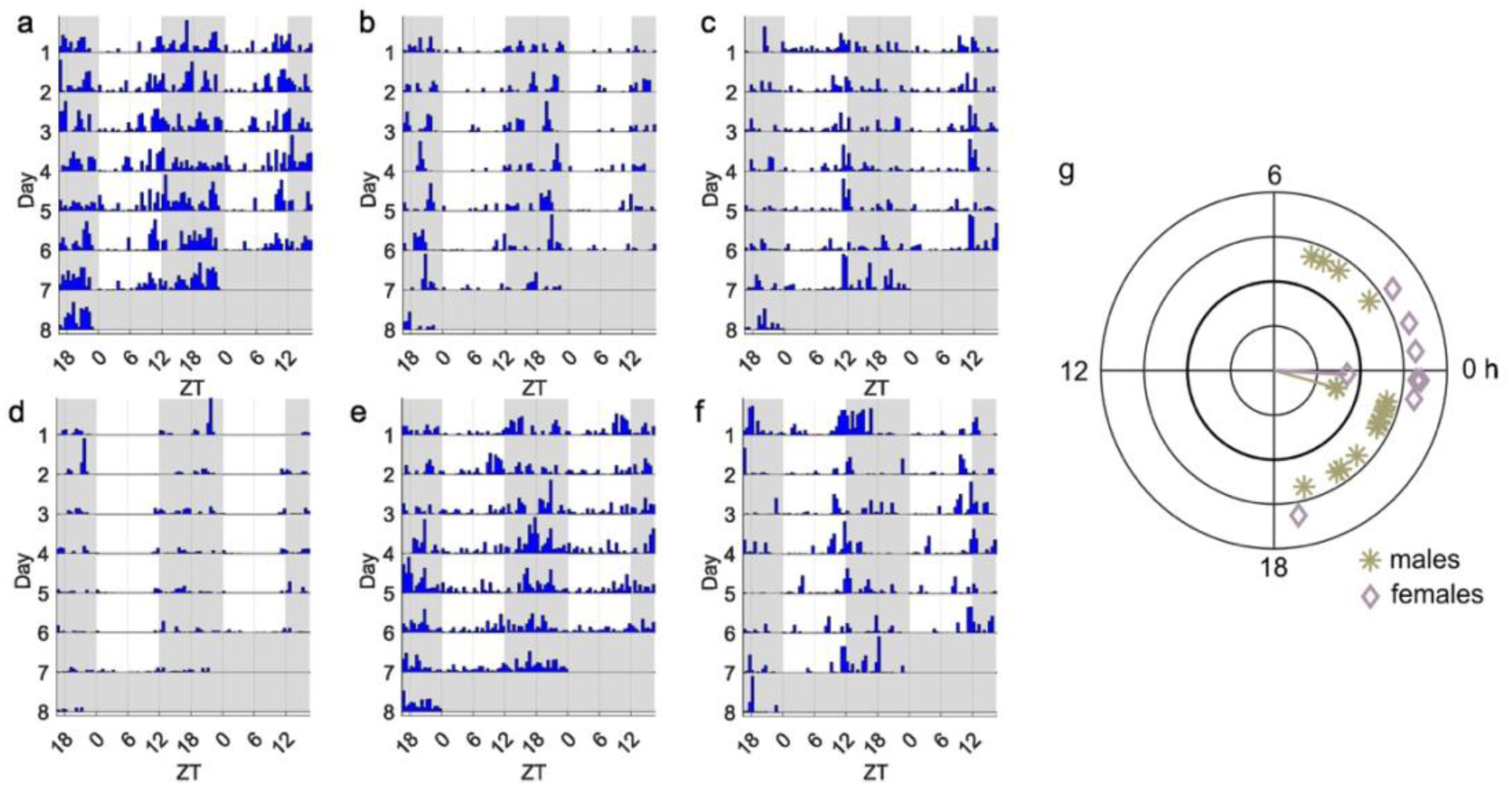
Variation among individuals. a-f Variation in behavioral patterns among individuals during LD; g: circular phase plot showing phases of beetles in LD based on data from the first 3 days. The lengths of the line from the centre represents the coherence of phases within a group.

**Figure 3:**
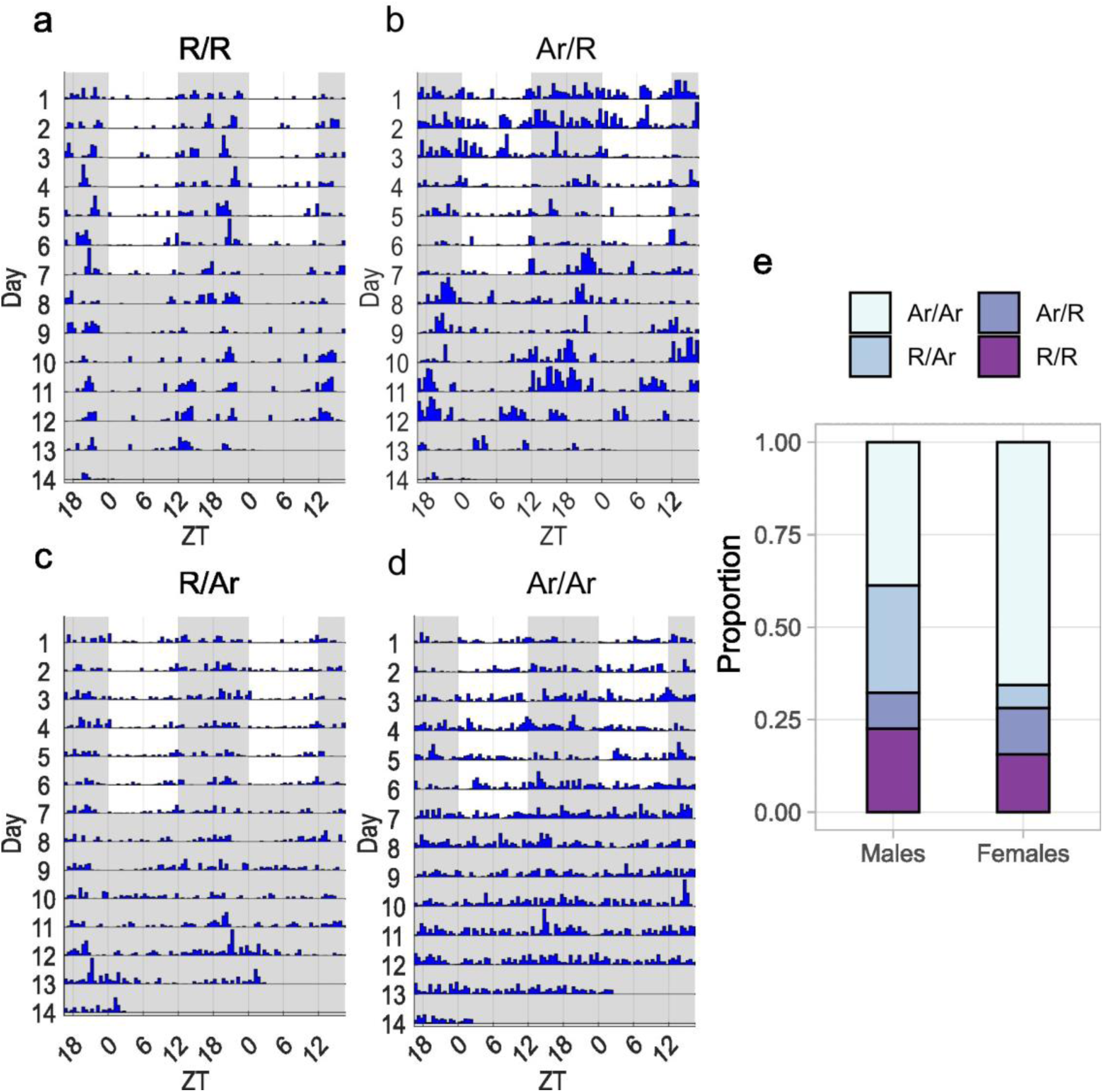
Maintenance of rhythmicity under LD and DD. a-d: Individual actograms showing different categories of beetles according to their status of being rhythmic (R) or arrhythmic (Ar) under the two conditions of LD (first 7 days) and DD (last 7 days) ( R/R - rhythmic in LD and DD, R/Ar – rhythmic in LD but not in DD, Ar/R – Rhythmic in DD but not in LD, Ar/Ar – Arrhythmic in both LD and DD), e: stacked bar plot showing the proportion of these categories separately for males and females.

### Locomotor activity patterns under constant light conditions

When LD was followed by constant light conditions (LL), the average activity profiles of beetles resembled those observed in DD, however, overall rhythmicity compared to DD was reduced under both light intensities tested (20 lux and 100 lux; Fig 4 and Fig S3, respectively, Table 1). When examining individual rhythmicity, males showed a significantly higher proportion of rhythmic individuals compared to females (GLMM, estimate = 1.711, p <0.001, Fig 4e). Furthermore, the proportion of rhythmic beetles experienced a significant decrease under constant light conditions compared to LD (GLMM, estimate = -3.193 p < 0.001, Fig 4e). There was an overall significant effect of light intensity on the rhythmicity (GLMM, estimate = -1.002, p = 0.03, Fig 4e). There was a significant interaction between light intensity and the light condition, with a sharper drop in rhythmicity from LD to LL for the higher light intensity (GLMM, estimate = 1.772, p= 0.026, Fig 4e).

**Figure 4:**
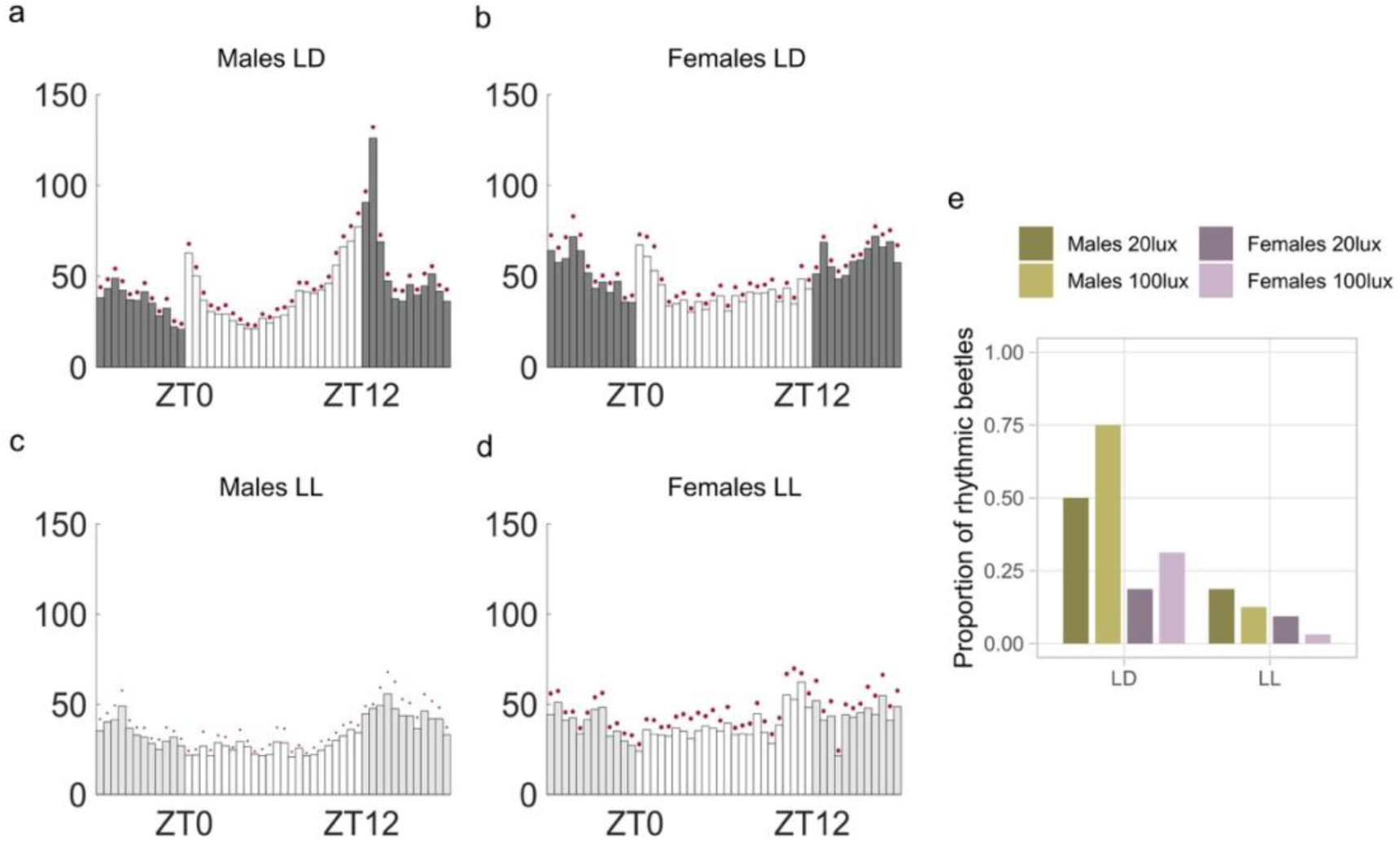
Rhythmicity under LL conditions. a-d Average histograms showing mean activity for all the beetles over the 7 days of recording under LD (light entrainment of 20 lux) and LL (20 lux) (Average histograms for 100 lux given in Fig S3), e: Proportion of rhythmic beetles in LD and LL for two light intensities;20 lux and 100 lux (males: n = 32 per light intensity, females: n=32 per light intensity)

### Population differences in locomotor activity and behavioral patterns

Both Cro1 and SB populations had similar behavioral patterns (Fig 5 a-h), but the average activity was significantly lower for the SB population during LD (GLMM, estimate = -0.667, p < 0.001, Fig 5 j) and DD (GLMM, estimate = -0.368, p = 0.04, Fig 5k). The average activity of females was significantly higher than that of males in LD (GLMM, estimate = -0.624, p < 0.001, Fig 5 j) but not in DD (GLMM, estimate = -0.118, p = 0.512, Fig 5k). While examining individual rhythmicity, the SB population showed a significantly higher proportion of rhythmic beetles in LD (GLMM, estimate = 1.116, p =0.006, Fig 5i). The effect of light condition was also significant with a higher proportion of beetles observed in LD (GLMM, estimate = 1.761, p <0.001, Fig 5 i). However, there was no significant effect of sex on rhythmicity (GLMM, estimate = 0.413, p = 0.279, Fig 5 i).

**Figure 5:**
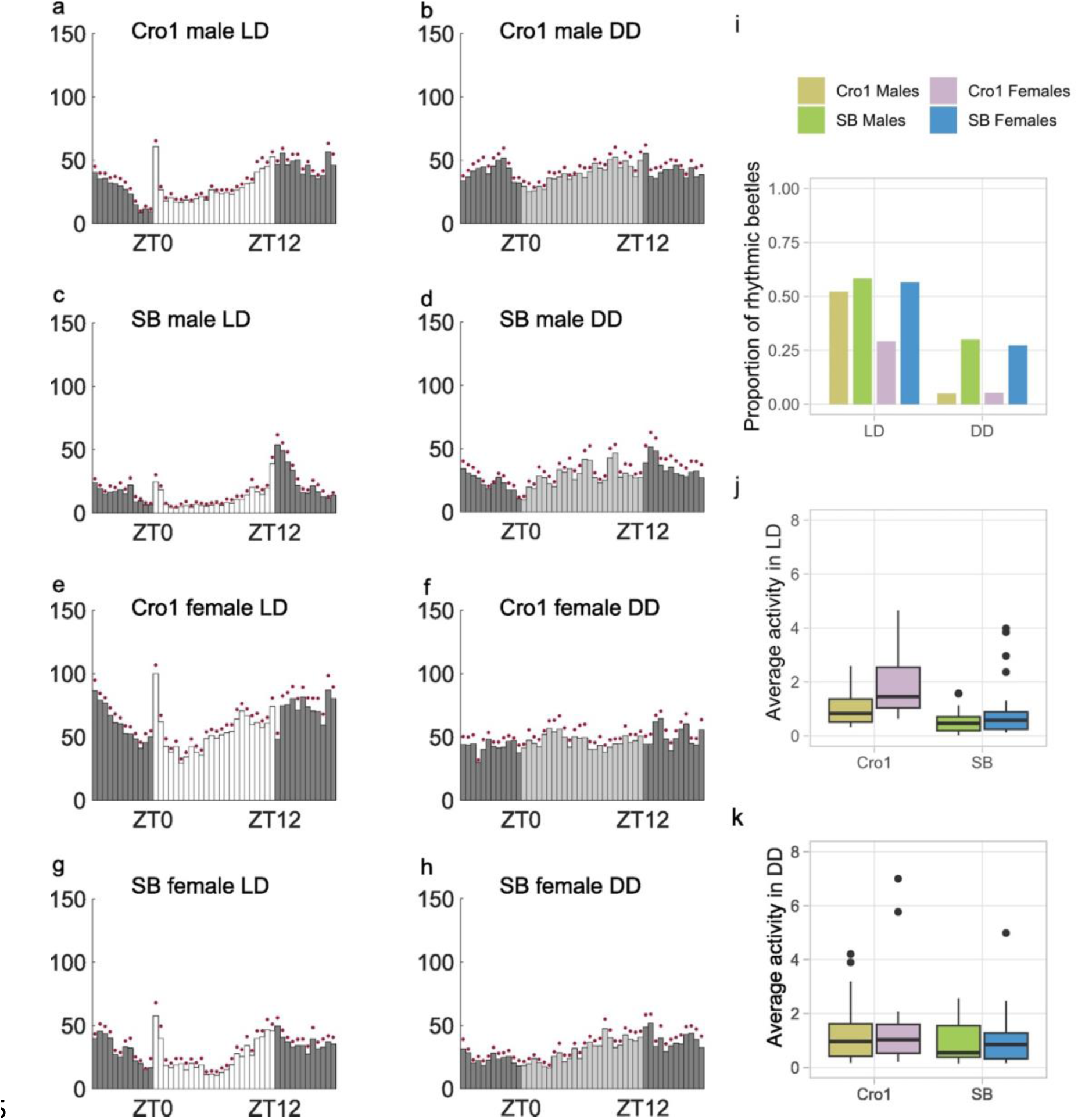
Population differences in rhythmicity. a-h: Average histograms showing mean activity for all the beetles over the 7 days of recording under LD (light entrainment of 20 lux) and DD for Cro1 and SB populations, i: Proportion of rhythmic beetles in LD (Cro1 males: n = 23 , Cro1 females: n=24, SB males: n=24, SB females: n=23) and DD (Cro1 males: n = 20 , Cro1 females: n=19, SB males: n=20, SB females: n=22) j and k: Boxplots showing activity in 1-minute bin for individual beetles averaged over 7 days in LD (j, Cro1males: n = 23, Cro1 females: n=24, SB males: n = 24, SB females: n = 23) and DD (k, Cro1 males: n = 22, Cro1 females: n=20, SB males: n = 21, SB females: n = 22)

### Locomotor activity patterns during and after temperature entrainment

Similar to the behavior in LD, the beetles showed activity rhythms in a 12 hr: 12 hr temperature cycle of 25^0^C: 30^0^C in constant darkness (DD) (Fig 6 a-d, Fig S4). However rhythmic behavioral patterns, apart from the prominent peaks of activity at the start of temperature change in the subjective morning and evening, were less prominent compared to LD. The peaks of activity at the temperature transitions resembled the startle response observed after lights-on in LD and were not retained in constant temperature conditions (Fig 6 a-d).

**Figure 6:**
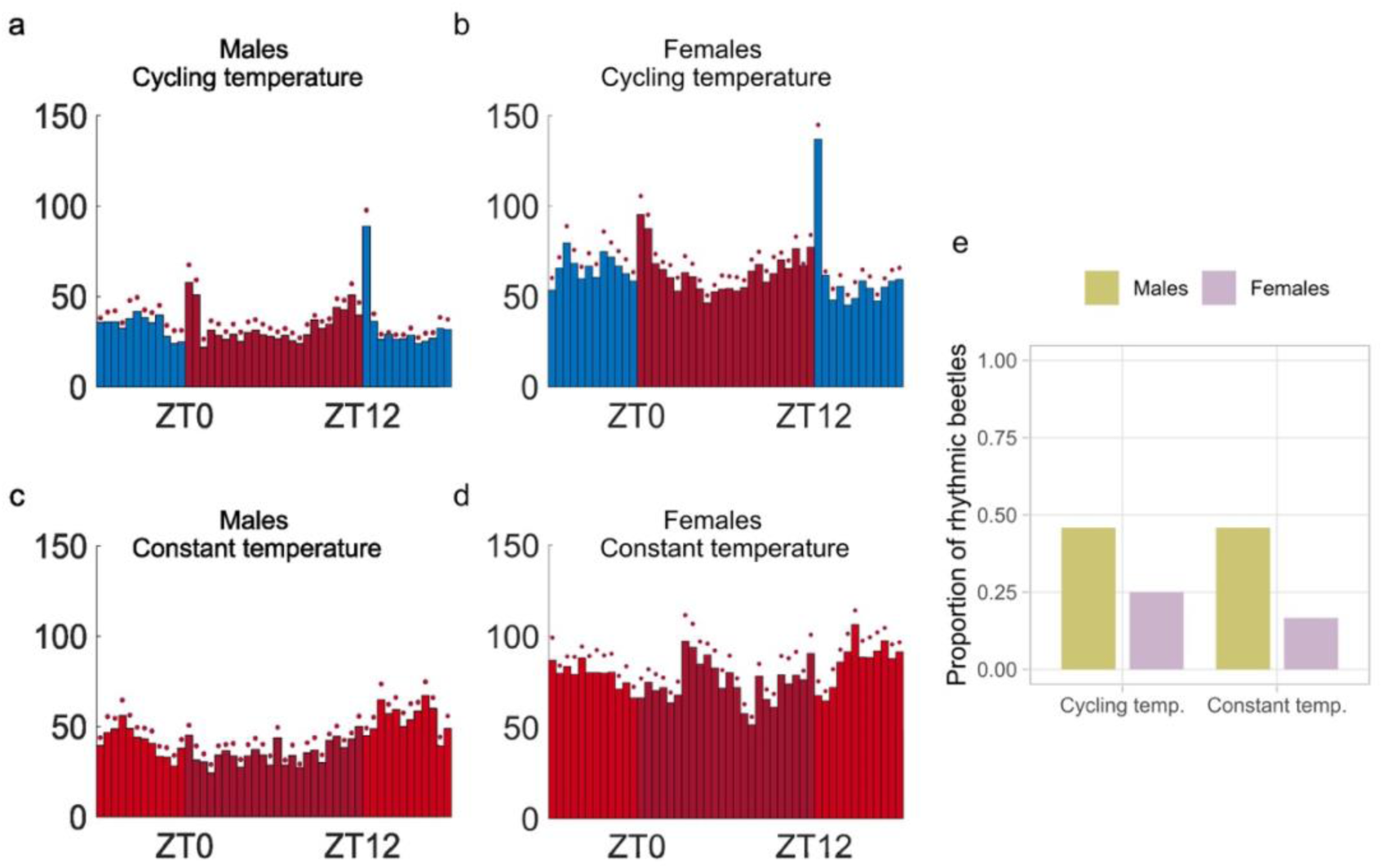
Temperature entrainment. a-d Average histograms showing mean activity for all the beetles over the 7 days of recording under temperature cycling and subsequent constant temperature, both under constant darkness, e: Proportion of rhythmic beetles in cycling temperature and constant temperature (males: n = 24 per temperature condition, females: n=24 per temperature condition)

After the temperature cycles, only males showed a clear rhythmic activity pattern under constant temperature conditions (Fig 6 c). The proportion of rhythmic beetles significantly differed between sexes (GLMM, estimate = 1.418, p = 0.018, Fig 6 e).

Upon the more natural-like temperature cycling in constant darkness with a ramping of temperature change within 2 hours, the activity patterns in the entrainment phase appeared stronger compared to the entrainment with a square-wave of temperature cycling (Fig S5). Both the startling peaks disappeared and there was a clear dip in the morning activity as well as a stronger evening anticipation in the entrainment phase resembling the pattern in LD (Fig S5). In the subsequent constant temperature conditions, both males and females retained these activity patterns. There was no difference in the proportion of rhythmic beetles between the sexes (GLMM, estimate = 0.185, p = 0.635, Fig S4 E) as well as between temperature cycles and constant temperature conditions (GLMM, estimate = -0.427, p = 0.283, Fig S4 e).

### Locomotor activity patterns upon knockdown of *Clk*

We downregulated the key circadian clock gene *Clk* using RNAi in the pupal stage to study the subsequent changes in activity profiles.

*Clk* gene expression was significantly different between the sexes (Table 2) in the RNAi control. There was a significant reduction in *Clk* gene expression compared to the RNAi control, indicating that dsRNA-mediated knockdown was efficient (Table 3). Following the *Clk* knockdown, there was also a significant reduction in rhythmicity (GLMM, estimate = -1.369, p = 0.005) in comparison to the RNAi control beetles, showing that *Clk* function is important for behavioral rhythmicity (Fig 7). There was a significant interaction effect of sex and light condition as well on the rhythmicity (GLMM, estimate = 1.391, p = 0.04) (Fig 7).

**Figure 7:**
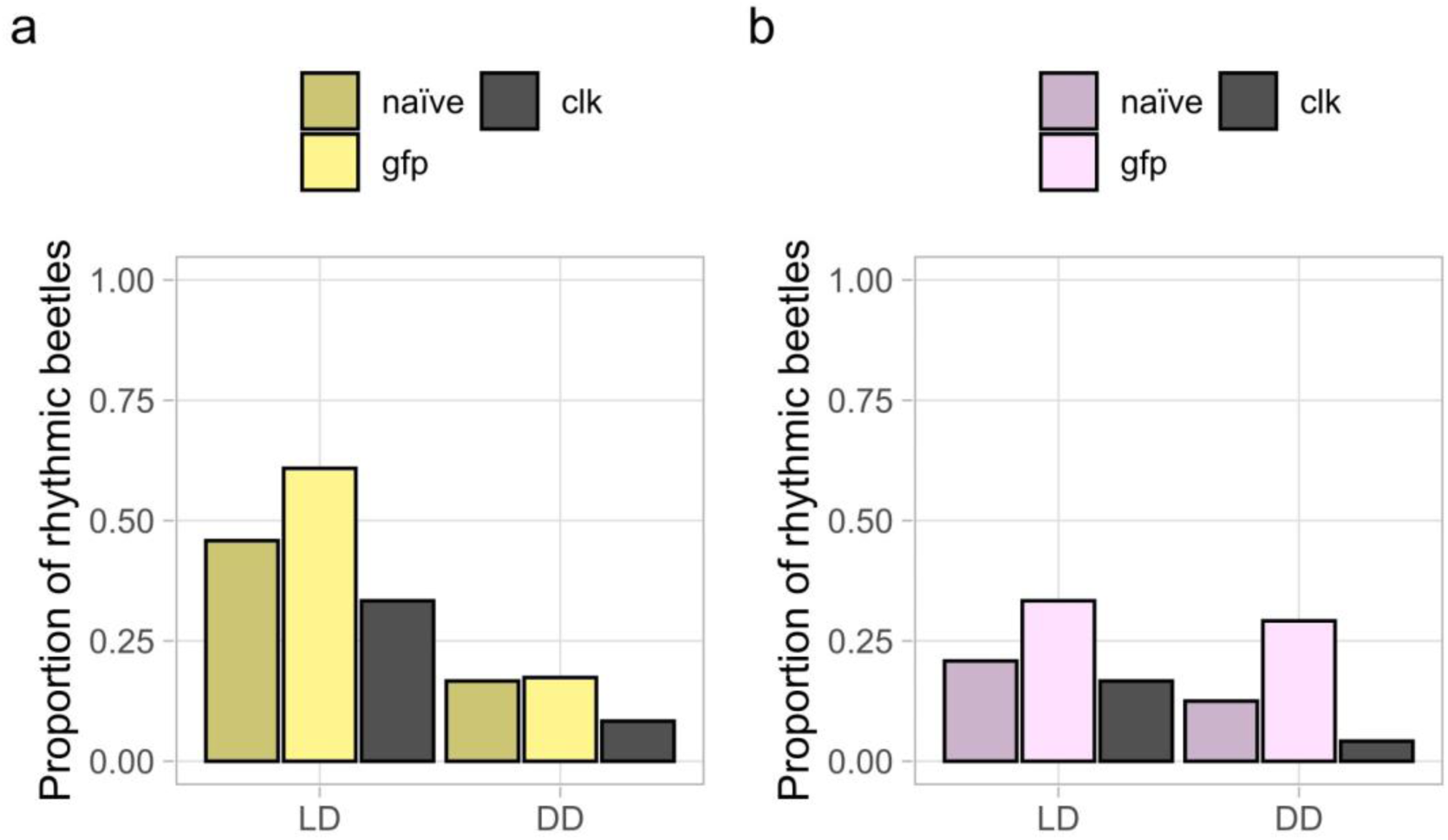
Rhythmicity upon knockdown of *Clk.* Proportion of rhythmic individuals in adults following the knockdown treatment at pupal stage for *Clk* in males (a) and females (b) (LD: n =24 per sex and treatment groups, DD: n=24 per sex and treatment groups except for RNAi control males (n = 23))

**Table 2:** Sex difference in gene expression of *Clk*

| Relative gene expression of males compared to females | 95% C.I. | n | p value |
| --- | --- | --- | --- |
| 0.312 | 0.158 - 0.795<br>ctrl vs ctrl (male vs male): 0.381 - 2.642 | 4 | **0.02** |

**Table 3:** Knockdown of *Clk*

| Sex | Relative expression compared to RNAi control | 95% C.I. | n | p value |
| --- | --- | --- | --- | --- |
| Male | 0.463 | 0.227 - 1.273<br>ctrl vs ctrl: 0.381 - 2.642 | 4 | **0.031** |
| Female | 0.531 | 0.279 - 0.988<br>ctrl vs ctrl: 0.539 - 1.858 | 4 | **0.028** |

## Discussion

We used the Drosophila Activity Monitoring system (DAM) to perform a detailed characterization of the average as well as individual locomotor activity profiles of adult *T. castaneum* extending its application to study behavioral patterns in another important model organism. We recorded the activity of lab-adapted populations with high temporal resolution and described the activity patterns under light entrainment, constant dark conditions and for the first time to our knowledge, under thermal oscillations and constant light conditions. The beetles anticipated environmental transitions and even though less robust, maintained their rhythmicity in DD and LL. We showed that a substantial amount of individual variation exists in the diel patterns of this cryptozoic species. The knockdown of *Clk* significantly reduced rhythmicity, strongly supporting the existence of circadian clock in *T. castaneum* and its control of behaviors. The finding that temperature cycling acts as a zeitgeber for the beetle clock, in addition to light, is consistent with our expectations, considering that beetles are exposed to cycling temperatures in their natural habitats of grain silos (Ziegler et al, 2021) as well as in their supposed original habitats under tree bark (Good, 1936; Sokoloff, 1974). Studies in many other insects including fruit flies (Glaser and Stanewsky, 2005) and honeybees (Giannoni-Guzmán et al., 2021) have documented the role of light-dark cycles as well as temperature oscillations as zeitgebers. The notable patterns of the average activity profiles under both zeitgebers included a consistent drop of activity just before the change of conditions in the morning which we refer to as ‘morning dip’ and the gradual increase of activity throughout the late afternoon known as evening anticipation. These patterns emerged even before the zeitgeber conditions change as well as appeared consistently in constant conditions, indicating an anticipatory response to the predicted changes resulting from their entrained molecular clock. Other insects such as *D. melanogaster* (Dubowy and Sehgal, 2017) and *Plutella xylostella* (D. Wang et al., 2021) also show such gradual increase in activity anticipating the transitions of environmental conditions. However, a gradual decrease of activity (morning dip) towards the time of dark-light transition is rather atypical.

The beetles reacted with a startle response of an exceptionally heightened activity to the light turning on in the morning as well as to the change of temperature in the subjective morning and evening. The absence of these startle responses in constant conditions as well as their disappearance or strong reduction under more natural-like conditions of ramping temperature or light conditions confirms that they are not mediated by the circadian clock but are instead reactions to sudden changes in environmental conditions. Similar startle responses immediately following the dark-light transition were observed in *Ae. aegypti*, and, consistent with our study, these responses were not present in constant darkness (Gentile et al., 2009) .

Average histograms consistently exhibited evening anticipation in constant conditions as well as following the entrainment phase for both light and temperature as zeitgebers. A steep evening anticipation was characteristic of the average activity profiles under light entrainment, especially for males. The evening anticipation aligns well with the time of day when mating rates of the *T.castaneum* reach their maximum (Z. Wang et al., 2023). In red flour beetles, males initiate the copulatory behavior and must mate multiple times to ensure fertilization success, especially because of the last male sperm precedence (Fedina and Lewis, 2008). An increased activity of males in search of mates could be a potential reason why males possess a strong and clear evening anticipation peak compared to the females. Similarly, in *P. xylostella*, males were shown to have better rhythmic features than females under constant darkness in low temperature conditions (D. Wang et al., 2021). Such sex-specific differences in activity profiles are commonly observed in many insect species and are often associated with their mating behavior. For example, in *Anopheles gambiae*, males have an earlier flight activity peak compared to females, consistent with the fact that males aggregate to form swarms to which females subsequently enter (Rund et al., 2012). In addition to the difference in evening anticipation, the overall activity levels of females were significantly higher than those of males, as was also reported recently (Benita et al., 2024). we also observe a large inter-individual variation in phase for the males compared to females. Additionally, males generally exhibit a higher proportion of rhythmicity, which may be attributed again to the necessity for males to initiate copulation behavior, thereby requiring precise timing of such behaviors.

In addition to the individual variation in phase, there was substantial variation in the individual activity patterns including differences in the duration of peak activity, levels of activity at different times, and the ability to maintain rhythmic behavior under constant conditions following entrainment. A study in the cave beetle species, *Laemostenus latialis* also showed similar variability in individual behavioral patterns (Pasquali and Sbordoni, 2014). Another study observed significant diversities in circadian behaviors in social insect species compared to the solitary species like fruit flies (Giannoni-Guzmán et al., 2014). Since *T.castaneum* is a non-social yet group living insect, one could speculate that the dual role of the pheromone, 4,8-dimethyldecanal (DMD)as an aggregation as well as a sex attractant for females (Levinson and Levinson, 1995) enables individuals to rely on their conspecifics for timing their own mating rather than a strict reliance on their circadian clock. Another possibility is that variability in their peak activity timings could help in avoiding any possible male-male competition.

The proportion of rhythmic beetles considerably decreased in DD conditions especially for males . The presence of all clock gene homologs known in *Drosophila* (Richards et al., 2008) indicates that *T. castaneum* possesses a functional circadian clock. Therefore, the decrease in rhythmicity in constant darkness compared to that under light entrainment could possibly indicate a weak link or uncoupling between the endogenous clock and the locomotor behavior under constant darkness (Abhilash et al., 2017) as seen in the case of blind mole rats (Beale et al., 2016) and in *Drosophila* species inhabiting higher latitudes with weakly rhythmic environments (Bertolini et al., 2019). The inter-individual variability seen in locomotor activity profiles could also possibly be the result of a weak behavioral regulation by the endogenous clock (Beale et al., 2016). One could speculate that such a weak coupling or even uncoupling might be due to the absence or reduced selective pressure for circadian functions in their possible ancestral natural dark habitats such as rotting logs and tree barks as well as in their recent, also largely dark habitats of grain silos (Good 1936; Sokoloff 1974). The possible chance of relying on olfactory cues from conspecifics for mating initiation might have also resulted in a weak selection pressure for the evolution of a strong endogenous clock or a strong coupling of the clock with behaviors.

There was a significant decrease in the proportion of rhythmic beetles in constant light conditions, more pronounced under higher light intensity, even if weak rhythmic patterns were observable in the average histograms. In wild-type D. *melanogaster*, strong constant light leads to arrhythmic behavior (Konopka et al., 1989, Matsumoto et al., 1994) because of the light-dependent TIM degradation by CRY1 which conveys photic signals to the circadian clock (Ceriani et al., 1999). The beetles are not expected to have such an arrhythmic behavior as they possess only the light-insensitive cryptochrome (CRY 2) (Yuan et al., 2007). However, most likely retinal and/or potentially also unknown extraretinal photoreceptors play a role (Helfrich-Förster, 2020) in photic entrainment, which could then possibly affect the rhythmicity under such constant light conditions. A similar reduction of rhythmicity under constant light conditions was reported for *Nasonia vitripennis*, which also possesses only the light-insensitive CRY 2 (Bertossa et al., 2014), whereas some other closely related *Nasonia* species retained their rhythmicity (Bertossa et al., 2013). A study in linden bug, which also possesses only CRY 2, revealed a deteriorating effect of constant light on rhythmicity even though around 40% of individuals showed strong rhythmicity under such constant light conditions (Kaniewska et al.,, 2020)

To test if the observed behavioral rhythms depend on a molecular circadian clock, we attempted the knockdown of the key circadian clock gene *Clk* in the pupal stage and subsequently monitored locomotor activity profiles at the adult stage. Indeed, rhythmicity in *Clk* dsRNAi animals was significantly reduced compared to the control *GFP* dsRNAi beetles, strongly supporting a role for a canonical CLK-dependent transcriptional/translational circadian feedback loop in *T. castaneum*. In the future, double knockdowns of different clock genes could be performed to induce more severe clock disruption and phenotypes.

Since *T.castaneum* is a non-social but gregarious species, further experiments to study their circadian rhythms in a group setting would be the next step. The social influence on circadian rhythms has not only been demonstrated in social insects, but also for a solitary species like *Drosophila* for which this is most likely mediated by olfactory cues (Levine et al., 2002a). Studies of circadian rhythms of *T.castaneum* in a social setting are especially important, because it is already known that an exposure to the male aggregation pheromone increases the expression of the *timeless* clock gene in these beetles (Rath et al., 2021).

As *T. castaneum* is a ubiquitous pest, knowledge about its diel patterns in behavior under different external environmental cues can be useful in pest management strategies, for instance in light traps. Recently, circadian clock research is gaining more importance in immune system homeostasis (Haspel et al., 2020). Improved knowledge about the circadian clock and the rhythms it controls in *T. castaneum,* which is an excellent model to study insect immunity (Altincicek et al., 2008; Ferro et al., 2019), can be beneficial for the emerging research in the intersection of circadian function and immune regulation which has potential clinical relevance. It is remarkable that despite the absence of strong light cues in their original habitats, the molecular clock of flour beetles still shows a strong entrainment to light as well as that many beetles can retain this rhythmicity under constant conditions. The considerable variation seen in different aspects of the diel patterns along with its status as a group living yet non-social insect, renders this organism a unique model for investigating circadian clocks and the different behavioral and physiological parameters they control. This complements studies in other insect groups, contributing to a comprehensive understanding of circadian rhythms across different insect species.

## Acknowledgments

We thank Hannah Goldgräbe for helping with the experiments.

## Note

Supplementary material for this article including the final data files is available online.

## Author contributions

JK and RS conceived the ideas. JK, RS, MO, RR, NS and TP planned the experimental procedure. RR, TP, NS and PM carried out the experiments and RR carried out the statistical analyses together with NS, MO, TP and PM. RR wrote the first draft which was revised by NS and TP. RR and TP made the figure panels. All authors contributed to writing of the article in further revisions and approved the submitted version.

