## Supplementary Materials for "Deciphering a beetle clock: individual and sex-dependent variation in daily activity patterns"

for the manuscript:

**Running title: Locomotor activity rhythms in *T. castaneum***

**Authors: Reshma R[1], Tobias Prüser[1], Nora K. E. Schulz[1], Paula M. F. Mayer[1], Maite Ogueta[2], Ralf Stanewsky[2,3#], Joachim Kurtz[1,3*#]**

[1]Institute for Evolution and Biodiversity, University of Münster, Hüfferstr. 1, 48149 Münster, Germany

[2]Institute of Neuro- and Behavioural Biology, University of Münster, 48149 Münster, Germany

[3] Joint Institute for Individualisation in a Changing Environment, University of Münster and Bielefeld University

[#]Shared senior author

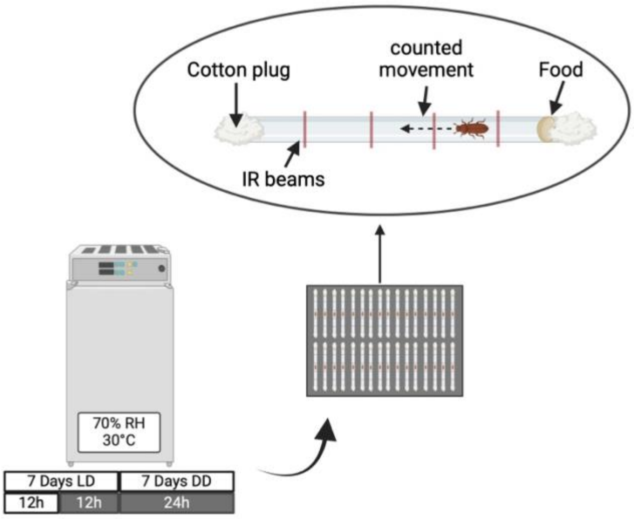


**Fig S1: Locomotor activity monitoring in Drosophila Activity Monitoring System (DAM5H-4)**. Individual beetles are placed in glass tubes with dried flour plate as food on one end and plugged with cotton on both ends. After arranging filled glass tubes into DAM5 systems, they are placed in an incubator. The monitor counts how often each beetle interrupts one of the four infrared light beams which penetrate the glass tubes.

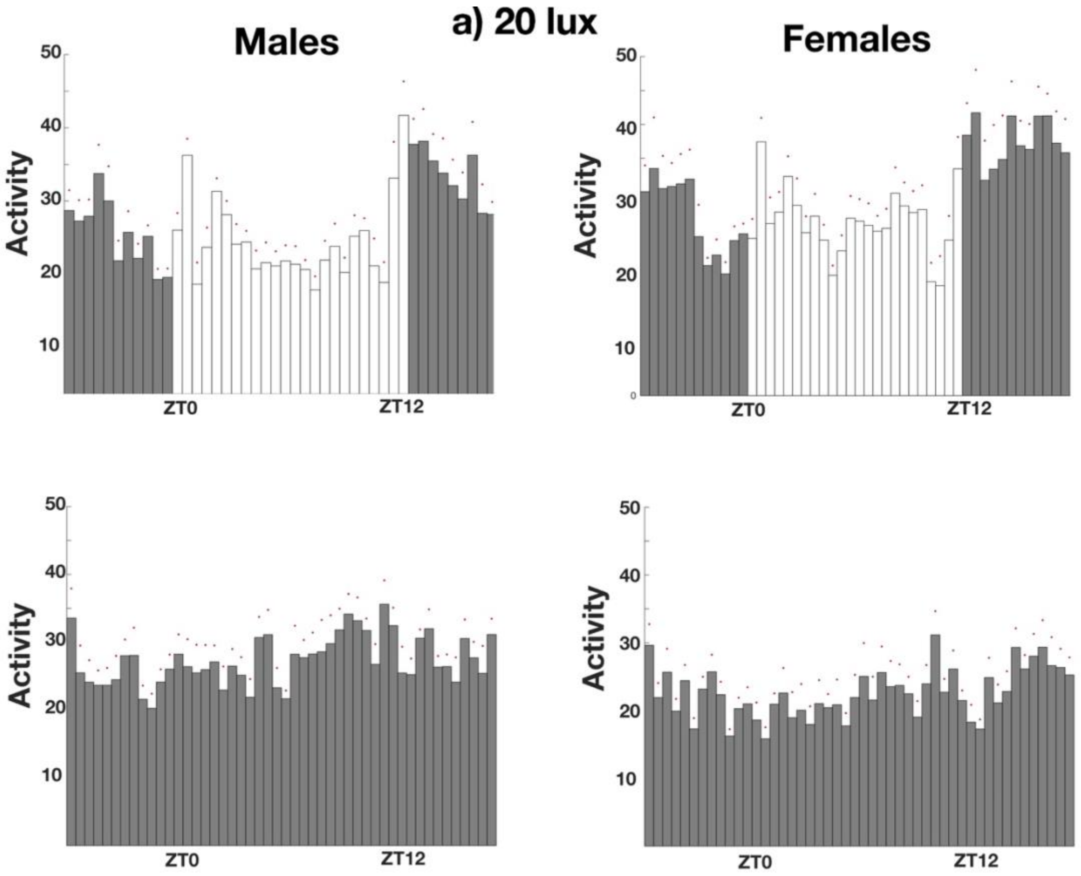
a) 20 lux
Males
Females
Activity
50
40
30
20
10
0
ZT0
ZT12

**b) 50 lux**

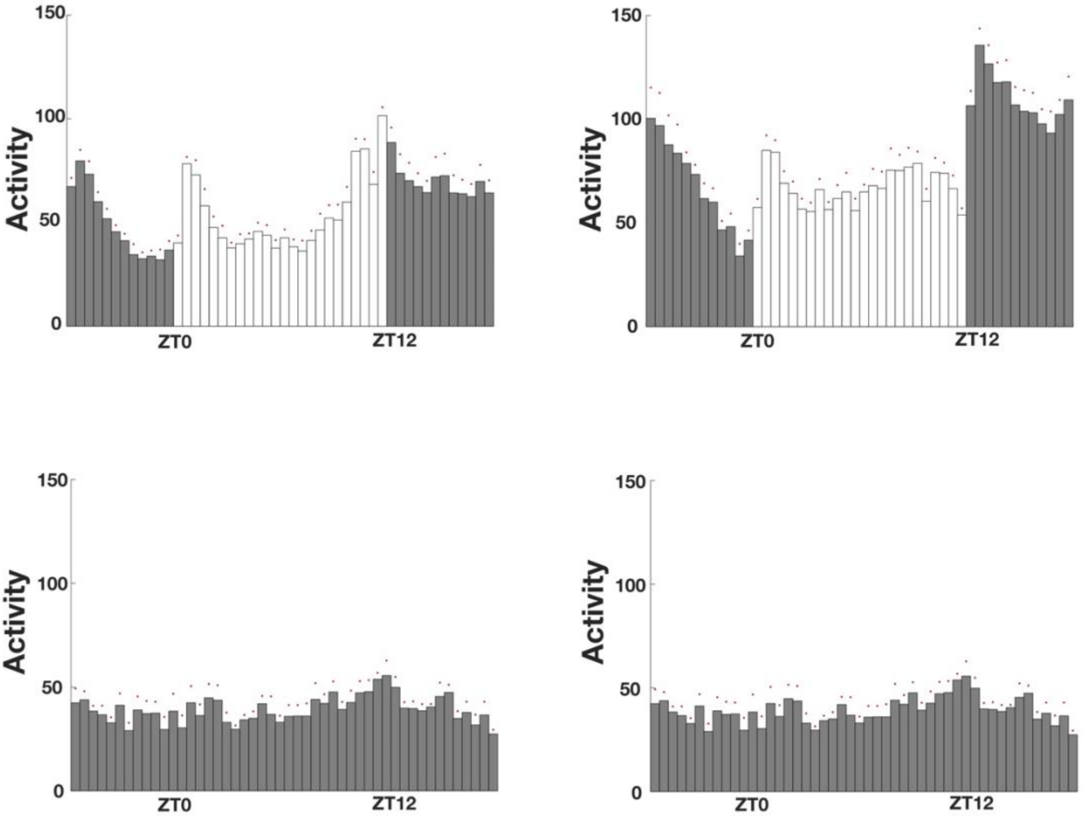
Activity
150
100
50
0
ZT0
ZT12
Activity
150
100
50
0
ZT0
ZT12
Activity
150
100
50
0
ZT0
ZT12
Activity
150
100
50
0
ZT0
ZT12

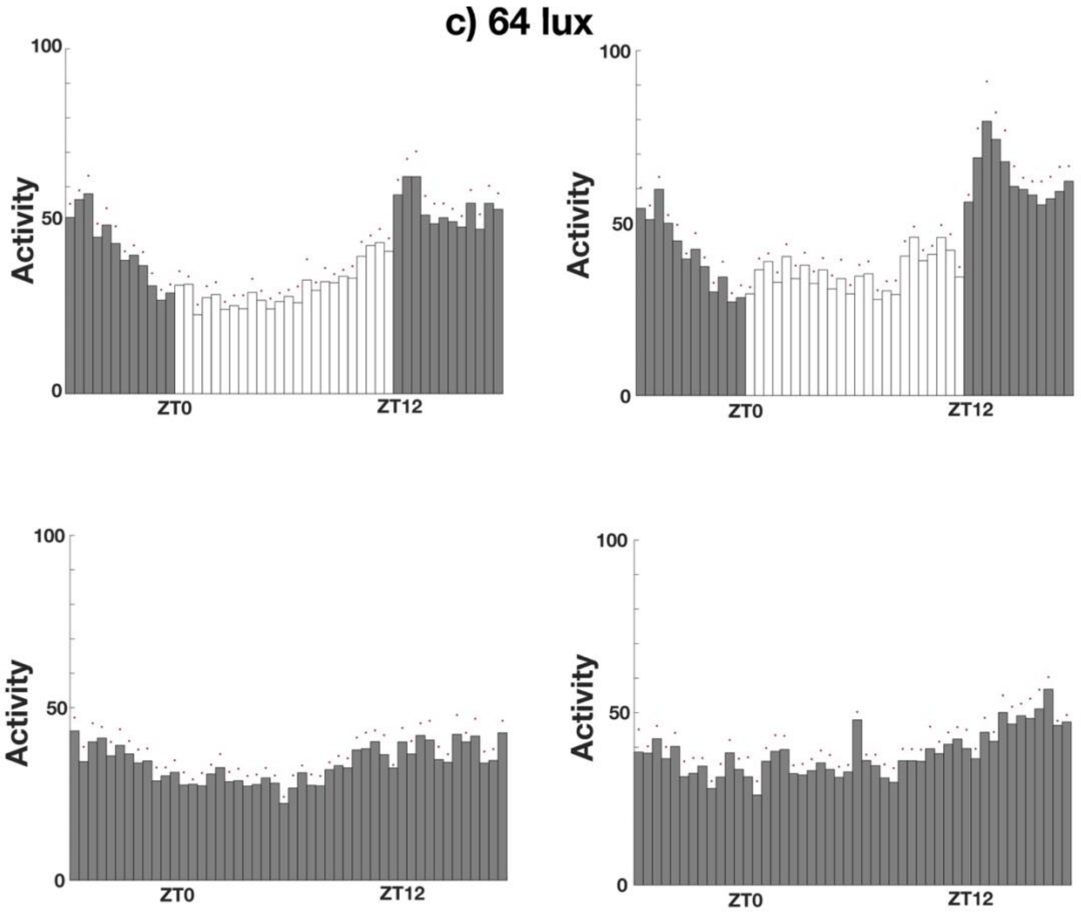


**Fig S2: Locomotor activity patterns of *T. castaneum* with light ramping**. Average histograms showing mean activity for all the beetles over the 7 days of recording under light entrainment with ramping followed by constant darkness with a lower (a- 20 lux: n = 31 per for males in each light condition, n = 32 in LD and n =31 in DD for females) and higher light intensities (b - 50 lux: n = 24 per sex for each light condition- 64 lux: n = 30 for males, n =32 for females)

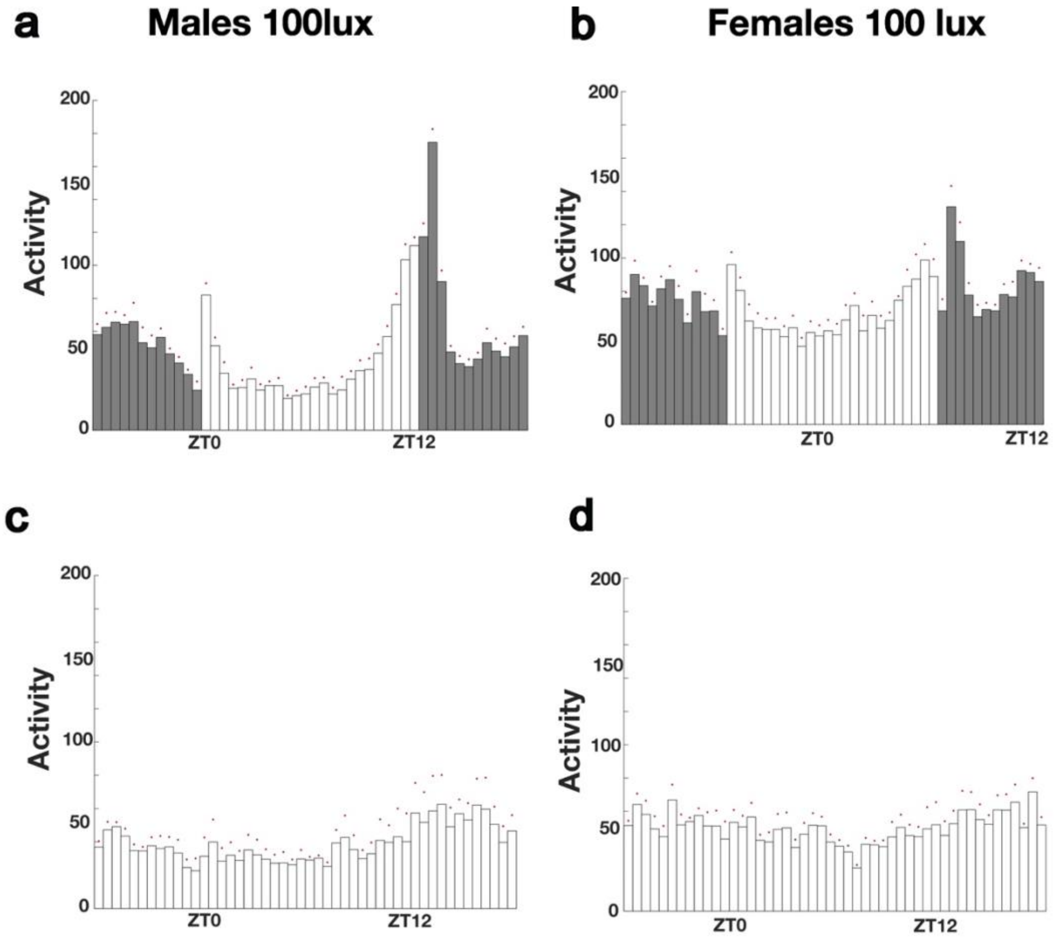


**Fig S 3 : a-d Rhythmicity under LL conditions (100 lux)**. Average histograms showing mean activity for all the beetles over the 7 days of recording under LD (light entrainment of 100 lux) and LL (males: n = 32 females: n=32)

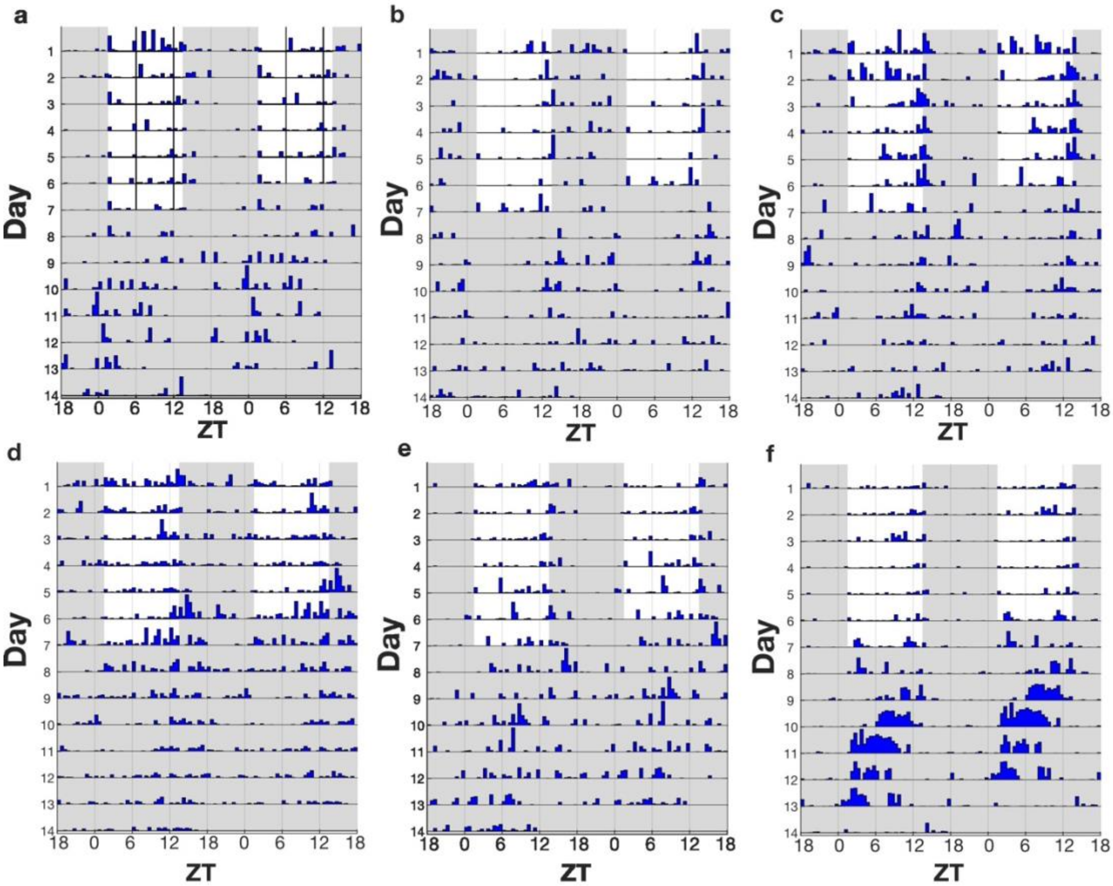


**Fig S4 Rhytmic beetles from temperature entrainment (without ramping).** Examples of individual beetles showing rhythmic behaviour in temperature entrainment followed by constant temperature with constant darkness maintained in both conditions.

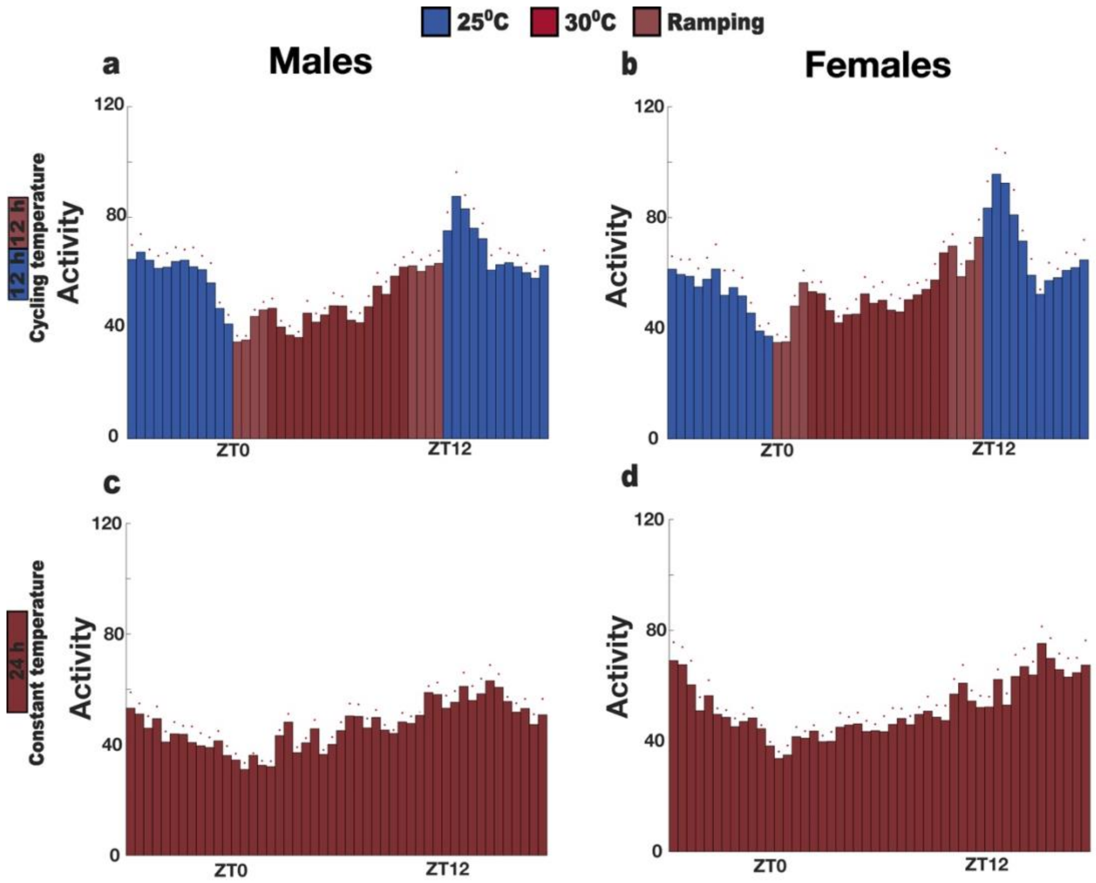
25⁰C
30⁰C
Ramping
a
Males
b
Females
12 h
12 h
Cycling temperature
Activity
120
80
40
0
ZT0
ZT12
c
d
24 h
Constant temperature

**e**

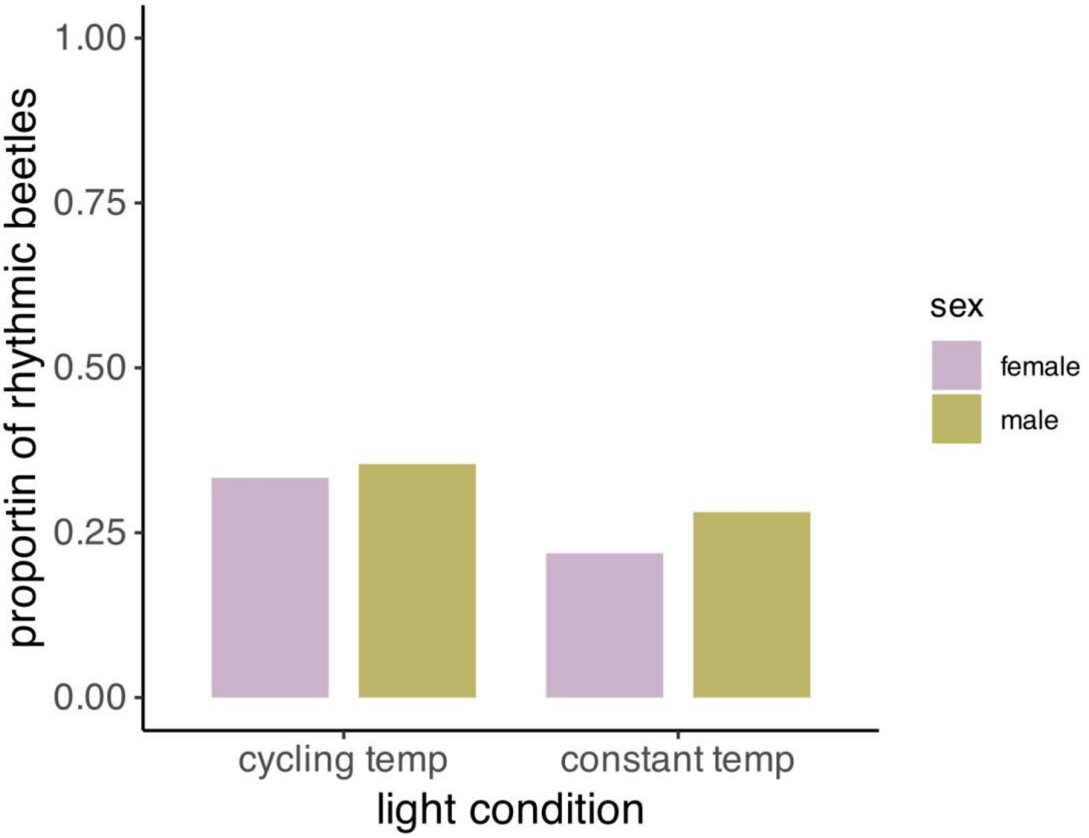


**Fig S 5:Temperature entrainment with ramping** a-d Average histograms showing mean activity in 30-minute bins for all the beetles over the 7 days of recording under temperature cycling with ramping followed by constant temperature both under constant darkness, e: Proportion of rhythmic beetles in cycling temperature and constant temperature (males: n = 48 in temperature cycling and n=32 in constant temperature, females: n = 48 in temperature cycling and n=32 in constant temperature)

**Table S1 :dsRNA used in this study**

| Gene name | NCBI Reference Sequence | iB number | Sequence |
|---|---|---|---|
| *Clock* (*clk*) | NM_001113466 /<br>TcasGA2_TC000088 | iB_00018 | TCCCGTGTTAACGAAATTCAAGAAGACT<br>GGAAACCGACGTTTCTCACAAACGAAG<br>AATTCACTCATTTAATCCTCGAAGCTGTG<br>GATGGCTTTATTATGGTGTTTTCGGC<br>TTCGGGTCAAATTTTTTATGCTTCGGAAA<br>GTATTACCTCGCTTTTGGGACATTTA<br>CCGAATCAAGTCCTGAATATGACAATATA<br>CGAAATGGCAAATGAAGAAGAACACA<br>GTCACCTCTACAATATTTTACTCACTCCGT<br>CCGAAGACCAGGGTCAAGTTTCCTT<br>CTCGTGTCATTTGCGACGTGGAGATCCT<br>GACTCCAAACAGAATCCCTCCTTTGAA<br>CTTGTGCATTTTGTAGGATATTTCAGATC<br>TGACGAAGATATGGTTCAGAGCGAAA<br>ATCG |
| *eGFP* | | | GTGAGCAAGGGCGAGGAGCTGTTCACC<br>GGGGTGGTGCCCATCCTGGTCGAGCTG<br>GACGGCGACGTAAACGGCCACAAGTTC<br>AGCGTGTCCGGCGAGGGCGAGGGCGAT<br>GCCACCTACGGCAAGCTGACCCTGAAGT<br>TCATCTGCACCACCGGCAAGCTGCCCGT<br>GCCCTGGCCCACCCTCGTGACCACCCTG<br>ACCTACGGCGTGCAGTGCTTCAGCCGCT<br>ACCCCGACCACATGAAGCAGCACGACTT<br>CTTCAAGTCCGCCATGCCCGAAGGCTAC<br>GTCCAGGAGCGCACCATCTTCTTCAAGG<br>ACGACGGCAACTACAAGACCCGCGCCG<br>AGGTGAAGTTCGAGGGCGACACCCTGG<br>TGAACCGCATCGAGCTGAAGGGCATCG<br>ACTTCAAGGAGGACGGCAACATCCTGG<br>GGCACAAGCTGGAGTACAACTACAACA |

| | | | |
|---|---|---|---|
| | | | GCCACAACGTCTATATCATGGCCGACAA GCAGAAGAACGGCATCAAGGTGAACTT CAAGATCCGCCACAACATCGAGGACGG CAGCGTGCAGCTCGCCGACCACTACCAG CAGAACACCCCCATCGGCGACGGCCCC GTGCTGCTGCCCGACAACCACTACCTGA GCACCCAGTCCGCCCTGAGCAAAGACC CCAACGAGAAGCGCGATCACATGGTCCT GCTGGAGTTCGTGACCGCCGCCGGGAT CACTCTCGGCATGGACGAGCTGTA |

**Table S2: Primers used in this study**

| Gene name | NCBI Reference Sequence | Sequence | Efficiency |
|---|---|---|---|
| *Clock* (*Clk*) | NM_001113466 / TcasGA2_TC000088 | F: ACTGCACCGAACACATGACA<br>R: ACGTGATAATTCCGCAACGC | 1.93 |
| Ribosomal protein L13a (*RpL13a)* | XM_969211.3 | F: GGCCGCAAGTTCTGTCAC<br>R: GGTGAA TGGAGCCACTTGTT | 1.98 |
| Ribosomal protein 49 (*Rp49)* | XM_964471.3 | F: TTATGGCAAACTCAAACGCAAC<br>R: GGTAGCATGTGCTTCGTTTTG | 1.95 |

**Table S3 Employed Generalized Linear Mixed Effect Models (GLMM) including distribution family**

| **Response variable tested** | **Final model** | **Family of error distribution** |
|---|---|---|
| Average activity levels in LD DD | average activity ~ light + sex +(1\|monitor) + (1\|ID) | gamma (link = "log") |
| Proportion of rhythmic beetles in LD DD | rhythmicity ~ sex*light + (1\|monitor) +(1\|ID) | binomial |
| Proportion of rhythmic beetles in LD LL | rhythmicity ~ sex + treatment*light + (1\|monitor) +(1\|ID) | binomial |
| Population differences in average activity in LD | average activity ~ population + sex + (1\|monitor) | gamma (link = "log") |
| Population differences in average activity in DD | average activity ~ population + sex + (1\|monitor) | gamma (link = "log") |
| Proportion of rhythmic beetles in temperature | rhythmicity ~ sex + temperature | binomial |

| | | |
|---|---|---|
| entrainment and constant temperature (no ramping) | condition+ (1\|monitor) +(1\|ID) | |
| Proportion of rhythmic beetles in temperature entrainment and constant temperature (with ramping) | rhythmicity ~sex +temperature condition + (1\|monitor) +(1\|ID) | binomial |
| Proportion of rhythmic beetles for knockdown of *Clk* | rhythmicity ~ sex + light + treatment + sex * light + (1 \| monitor) + (1\|ID) | binomial |
